# MHC class II in dopaminergic neurons prunes GABAergic synapses in neurodevelopmental disorders

**DOI:** 10.64898/2026.08.26.747425

**Authors:** Gen Murakami, Masataka Hirasaki, Miki Hashizume, Ayumi Hirao, Rina Ito, Yasushi Hojo, Takanari Nakano, Naonori Uozumi, Takayuki Murakoshi

**Affiliations:** Department of Liberal Arts, Faculty of Medicine, Saitama Medical University, 38 Morohongo, Moroyama-machi, Iruma-gun, Saitama 350-0495, Japan; Department of Clinical Cancer Genomics, Saitama Medical University International Medical Center, Yamane Hidaka, Saitama, 350-1241, Japan; Department of Biochemistry, Faculty of Medicine, Saitama Medical University, 38 Morohongo, Moroyama-machi, Iruma-gun, Saitama 350-0495, Japan

**Author notes:** Corresponding authors: Gen Murakami, Assistant Professor, Department of Liberal Arts, Faculty of Medicine, Saitama Medical University, Saitama 350-0495, Japan.

**Keywords:** mhc, major histocompatibility complex, neurodevelopmental disorders, maternal immune activation, MIA, dopaminergic neurons, glutamate decarboxylase, gad

## Abstract

Although the brain was traditionally considered immune-privileged, recent studies show immune factors play key roles in brain function. Dysfunction of these factors is linked to neurodevelopmental disorders, but mechanisms remain unclear. Using a maternal immune activation (MIA) mouse model, we investigated immune-related genes in neurodevelopmental disorder pathogenesis. MIA mice showed increased locomotor activity and disrupted prepulse inhibition. RNA-seq and qPCR analyses revealed persistent increases in major histocompatibility complex class II (MHCII) expression and persistent decreases in GABAergic synapse-related gene expression, particularly glutamate decarboxylase (Gad) expression, in dopaminergic regions. These expressions were negatively correlated, and immunohistochemistry showed MHCII at postsynaptic GABAergic synapses on dopaminergic neurons. Patch-clamp recordings confirmed reduced mIPSC frequency in MIA mice. MHCII knockout mice showed opposite phenotypes, while MHCII overexpression in dopaminergic neurons decreased Gad expression. These results suggest MIA-induced MHCII upregulation enhances pruning of GABAergic synapses on dopaminergic neurons, leading to behavioral deficits.

## Introduction

Although the brain has traditionally been considered an immune-privileged organ, recent studies have demonstrated that various immune factors, such as the complement system and microglia within the brain parenchyma, as well as T cells and monocytes at the brain borders, play important roles in the central nervous system ^1–4^. Furthermore, their dysfunction is associated with the pathogenesis of neurological disorders, including neurodevelopmental disorders such as attention-deficit hyperactivity disorder (ADHD), autism and schizophrenia ^5^. One of these immune factors is major histocompatibility complex (MHC) ^6^, and clinical studies have revealed significant associations of MHC with neurodevelopmental disorders ^7,8^. MHC refers to a group of cell surface proteins responsible for antigen presentation to immune cells, and predominantly consists of class I and class II, which are both classified into two categories, classical and nonclassical. MHC class I (MHCI) is expressed throughout the developing and mature brain and directly involved in the regulation of synaptic pruning ^9^. Our previous studies have also demonstrated that classical MHCI is expressed throughout the dopamine system and that its dysfunction underlies the pathogenesis of ADHD-like behaviors ^10–12^. On the other hand, the presence and function of MHC class II (MHCII) in the healthy brain remains controversial.

Maternal immune activation (MIA) is one of the most frequently used animal models for the study of neurodevelopmental disorders. This model mimics the environmental risk factor of immune activation during the fetal period by administering immune activating agents, specifically poly I:C, to pregnant animals. In this model, persistent epigenetic changes in immune pathways are considered critical for mediating these effects ^13,14^. Therefore, to comprehensively understand the pathogenesis of neurodevelopmental disorders, we investigated immune-related genes whose expression levels remained altered into adulthood in the brains of MIA-treated mice.

## Results

### MIA treatment-induced behavioral deficits relevant to ADHD and schizophrenia

In maternal immune activation (MIA) models, the type of drug, timing, and dose of treatment can result in different behavioral outcomes in adulthood ^14^. Therefore, we first evaluated our MIA treatment regimen (Figure 1A). Our treatment was sufficient to induce immune activation in the fetal brain three hours after treatment, as indicated by enhanced expression of interferon-gamma (*Ifng*), interleukin-6 (*Il-6*), and tumor necrosis factor-alpha (*Tnfa*) (Figure 1B).

**Figure 1.**
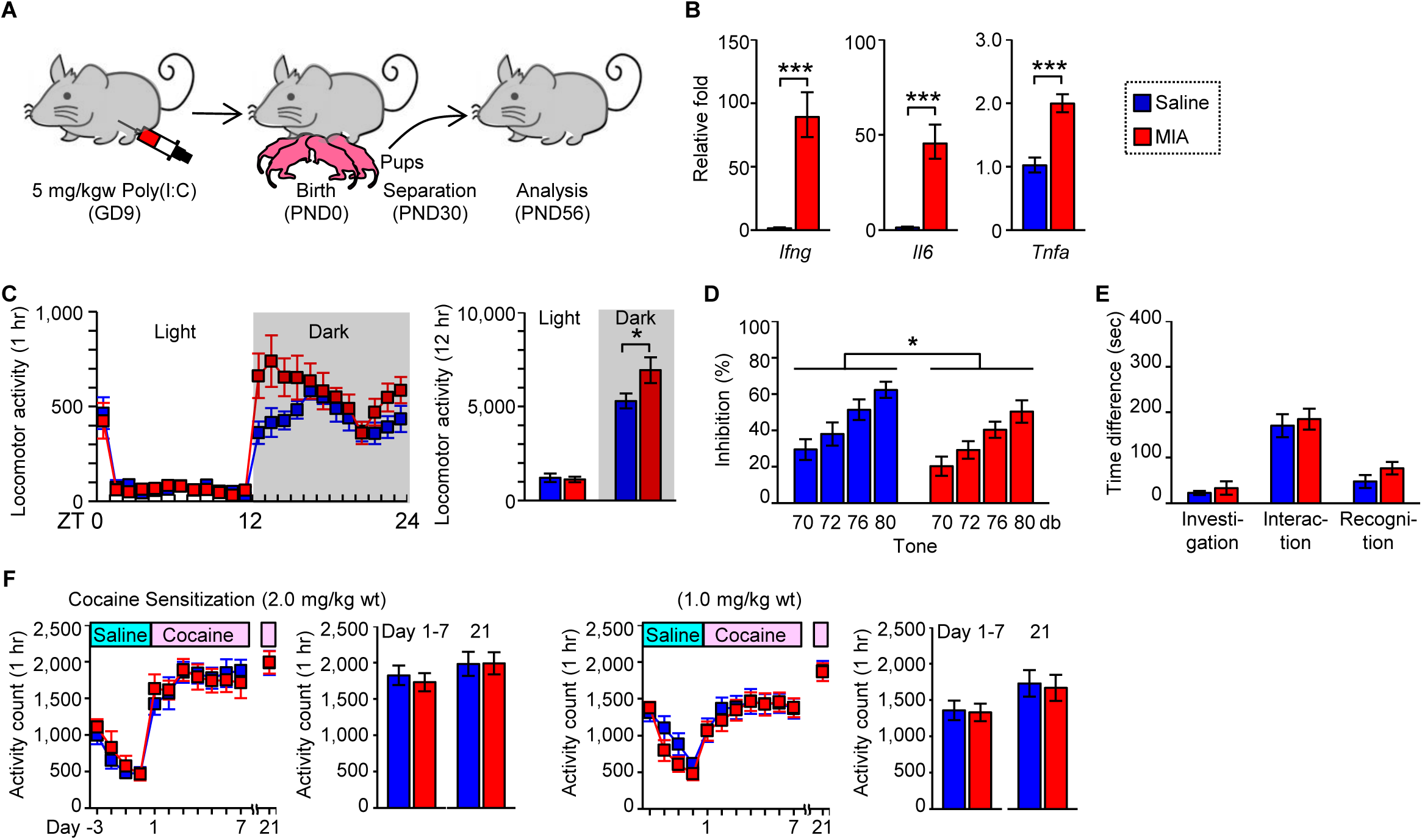
Behavioral deficits characteristic of neurodevelopmental disorders induced by MIA treatment. (A) Schematic diagram of the experimental paradigm. (B) Expression of proinflammatory cytokines in the whole-brain of fetal mice 3 hours after treatment with saline (*n* = 4) or MIA (*n* = 4) (Student’s t test, \*\**p* < 0.01). (C) Locomotor activity of mice treated with saline (*n* = 13) or MIA (*n* = 10) (Student’s t-test, \**p* < 0.05). (D) PPI ratio calculated by the response to a startle stimulus (120 dB) with a weak pre-stimulus (70, 72, 76 and 80 dB) divided by the startle stimulus only in mice treated with saline (*n* = 12) or MIA (*n* = 10) (two-way ANOVA, stimulus: F_(1, 80)_ = 13, \*\*\**p* < 0.001; treatment: F_(1, 80)_ = 7.0, *p < 0.05; stimulus × treatment: F_(3, 80)_ = 0.040, p = 0.99). (E) Investigation (left), social interaction (middle) and social recognition (right) in the three-chamber test compared between mice treated with saline (*n* = 13) or MIA (*n* = 10). (F) Locomotor activity in mice treated with saline (*n* = 8) or MIA (*n* = 8) received daily injections of cocaine (2.0 or 1.5 mg/kg body weight) for 7 days. After 10 days without injections, all mice were challenged with cocaine at the same dose as their previous injections. Locomotor activity was measured over 60 min after each injection. All data are represented as mean ± SEM and labeled as in (B).

We then evaluated locomotor activities, PPI, and social behaviors, which are characteristically impaired in ADHD, schizophrenia, and autism, respectively after the male offspring reached more than 9 weeks of age after MIA treatment. MIA-treated mice showed increased locomotor activity during the dark phase but not during the light phase (Figure 1C). MIA treatment also reduced PPI; however, this treatment had no significant effect on social behaviors (Figure 1D and 1E).

We further examined the effect of MIA on cocaine-induced behavioral sensitization, as neurodevelopmental disorders are significantly associated with drug addiction ^15^. Our previous studies have also demonstrated that MHCI deficiency induces both ADHD-like behaviors and enhanced cocaine-induced behavioral sensitization in mice ^11,12^. However, there were no significant differences between MIA- and saline-treated mice in cocaine-induced behavioral sensitization at either dose of 1.0 or 2.0 mg/kg body weight (Figure 1F). These findings indicate that our MIA treatment induces behavioral deficits relevant to ADHD and schizophrenia.

### Comprehensive analysis of MIA treatment-induced alterations in gene expression

Because long-lasting epigenetic changes in immune pathways are considered important for the transduction of MIA effects, we searched for immune-related genes whose expression levels changed up to adulthood after MIA treatment. We particularly focused on the dopamine system because this system is critically involved in neurodevelopmental disorders and particularly sensitive to proinflammatory cytokines ^16^. For this purpose, we conducted an exploratory RNA-seq analysis with a limited number of biological replicates for a comprehensive analysis of alterations in gene expression, using adult brain regions associated with the dopamine system of MIA-treated mice. These regions included the VTA, from which dopaminergic neurons originate, as well as the mPFC and NAc, which receive dense projections from dopaminergic neurons. Additionally, the Hip was included as a control region. We observed alterations in gene expression induced by MIA treatment in all examined brain regions. The greatest number of genes that were significantly altered with more than a two-fold change was observed in the VTA (Figure 2A).

**Figure 2.**
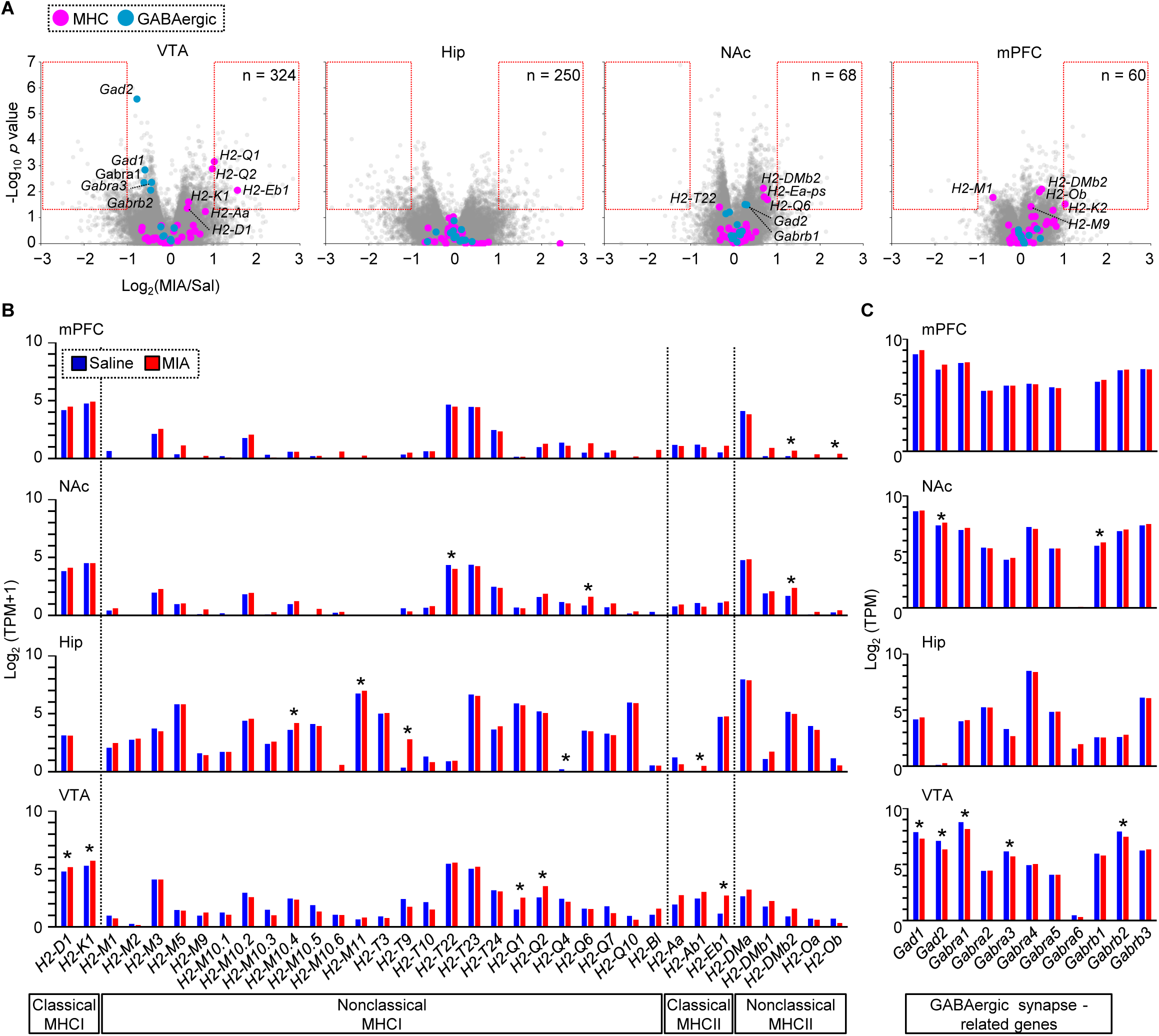
Comprehensive analysis of MIA treatment-induced alterations in gene expression by RNA-seq. (A) Volcano plot of RNA-seq analysis in the VTA, mPFC, NAc, and Hip of mice treated with saline (*n* = 3) or MIA (*n* = 3). MHC and GABAergic synapse-related genes are represented by purple and cyan circles, respectively. Genes within the red dashed rectangles indicate significantly altered genes, with more than a 2-fold change, and their numbers are shown in the upper right corner. (B and C) Comparative analysis of MHC (B) and GABAergic synapse-related gene expression (C) in the VTA, mPFC, NAc, and Hip of mice treated with saline (*n* = 3) or MIA (*n* = 3). All data are represented as mean ± SEM. \**p* < 0.05 and labeled as in (B).

In the analysis of immune-related genes in the VTA, MIA-treated mice exhibited significantly increased expression of classical MHCI genes (*H2-K1* and *H2-D1*), as well as nonclassical MHCI genes (*H2-Q1* and *H2-Q2*) (Figure 2B). These mice also exhibited significantly elevated expression of a classical MHCII gene (*H2-Eb1*), as well as a positive but not statistically significant increase in expression of the other classical MHCII genes (*H2-Aa* and *H2-Ab1*) and certain nonclassical MHCII genes (*H2-DM*). In contrast to MHC genes, the expression of nearly all complement system-, microglia-, or T cell-related genes remained unaltered by MIA treatment across all examined brain regions (Figure S1A). Next, we investigated genes that are specifically expressed in each type of synapse. MIA-treated mice exhibited decreased expression of genes related to GABAergic synapses, such as *Gad* or *Gabar* genes, particularly in the VTA, whereas much fewer genes related to glutamatergic synapses were altered (Figure 2C and S1B).

We also identified altered expression of dopamine receptor D5 (*Drd5*) and synapsin 1 (*Syn1*), both of which are associated with GABAergic synapses, as evidenced by the direct binding of DRD5 with GABA receptors and specific expression of SYN1 but not SYN2 or SYN3 at presynaptic sites of GABAergic synapses ^17,18^ (Figure S2A). In other brain regions, altered expression of classical MHC or GABAergic-related genes was rare or absent. These results suggest that classical MHC and GABAergic synapse-related genes in the VTA are possible candidates underlying MIA treatment-induced behavioral abnormalities.

### MIA treatment-induced increases in MHCII expression and decreases in *Gad* expression

To confirm the alterations in gene expression identified by RNA-seq in the VTA, qPCR analysis was conducted. We did not observe significant increases in the expression of all MHCI genes, but we did observe increases in the expression of all classical MHCII genes in MIA-treated mice (Figure 3A). Regarding genes related to GABAergic synapses, decreased expression of *Gad1/2*, but not GABA receptors, was confirmed in MIA-treated mice. This expression pattern of GABAergic synapse-related genes was also confirmed at the protein level (Figure 3B). These findings suggest that MIA treatment particularly affects expression of classical MHCII and *Gad* genes.

**Figure 3.**
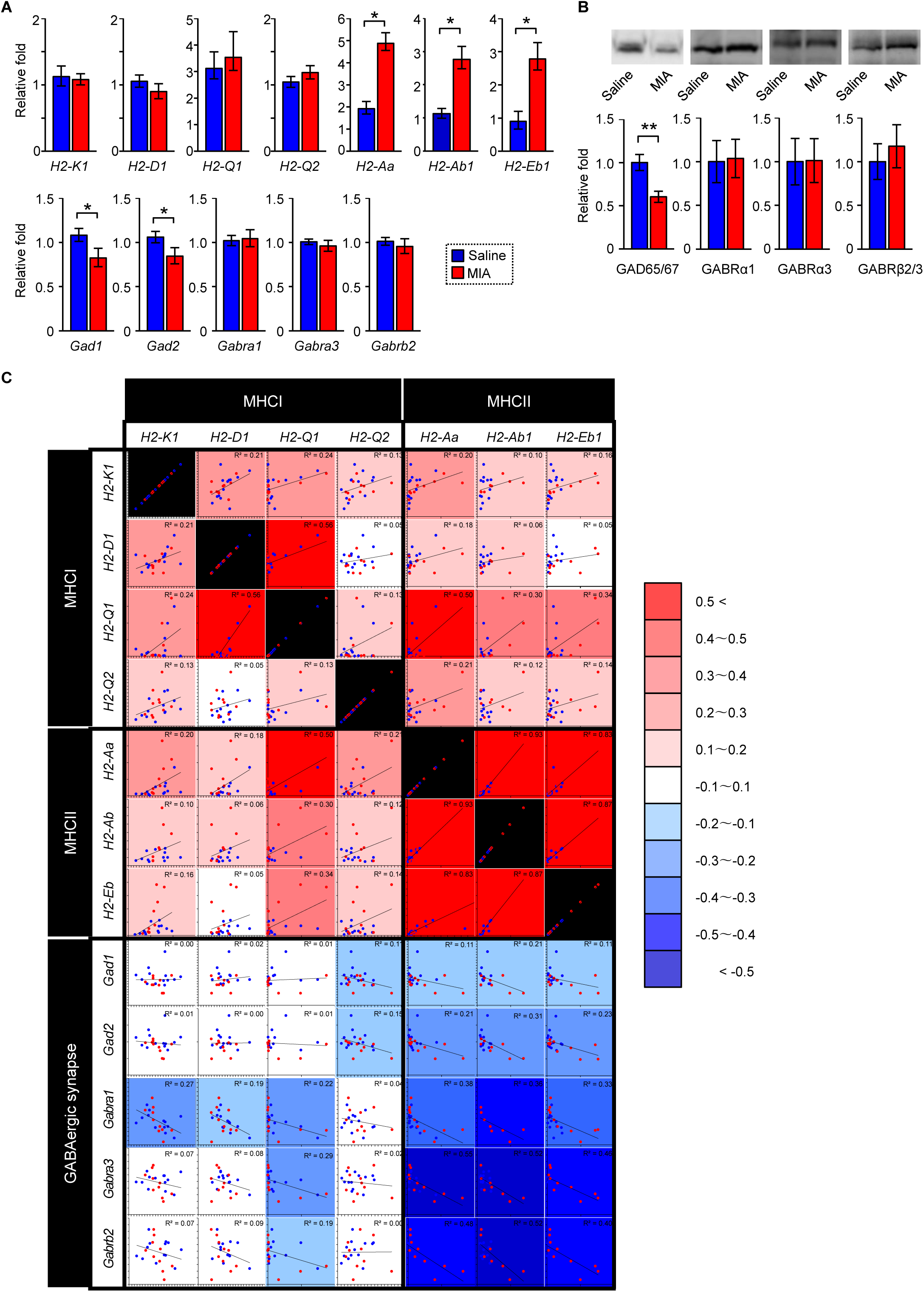
Comparative analysis of MIA treatment-induced alterations in gene expression by qPCR. (A) MHC and GABAergic synapses-related mRNA expression in the VTA of mice treated with saline (*n* = 13) or MIA (*n* = 10) (Student’s t test, \**p* < 0.05, and \*\**p* < 0.01). (B) GABAergic synapses-related protein expression in the VTA of mice treated with saline (*n* = 5) or MIA (*n* = 5) (Student’s t test, \*\**p* < 0.01). (C) Correlation analysis of MHC and GABAergic synapses-related mRNA expression in the VTA of mice treated with saline (*n* = 13) or MIA (*n* = 10). *r*^2^values are shown in the upper right corners. All data are represented as mean ± SEM and labeled as in (A).

Next, we conducted a correlation analysis, which has the advantage of evaluating associations between genes by accounting for individual differences among animals, making it more sensitive for detecting gene relationships than simply comparing group means after treatment ^19,20^. We found strong positive correlations among classical MHCII expression, as well as strong negative correlations of all classical MHCII with GABAergic synapse-related gene expression (Figure 3C). This analysis also revealed positive correlations of all classical MHCII with *Drd5*, but not with *Drd2* or *Dat,* as well as negative correlations with *Syn1*, but not with *Syn2* or *Syn3* (Figure S2B). These results demonstrate the specific association of MHCII with GABAergic synapses.

### MHCII localization at postsynaptic sites of GABAergic synapses

Because results from both RNA-seq and qPCR analyses revealed MHCII expression in the VTA, we sought to determine which cell types express MHCII in the VTA using immunohistochemistry. Based on our previous methodology, we first characterized the specificity of antibodies against MHCII in VTA slices prepared from WT and MHCII knockout (KO) mice which constitutively lack all classical MHCII genes (*H2^dlAb^*^1^*^-Ea^*) ^21^. Because a monoclonal antibody against MHCII (clone M5/114.15.2) showed specific staining in WT but not in KO mice, we used this antibody for our study (Figure 4A).

**Figure 4.**
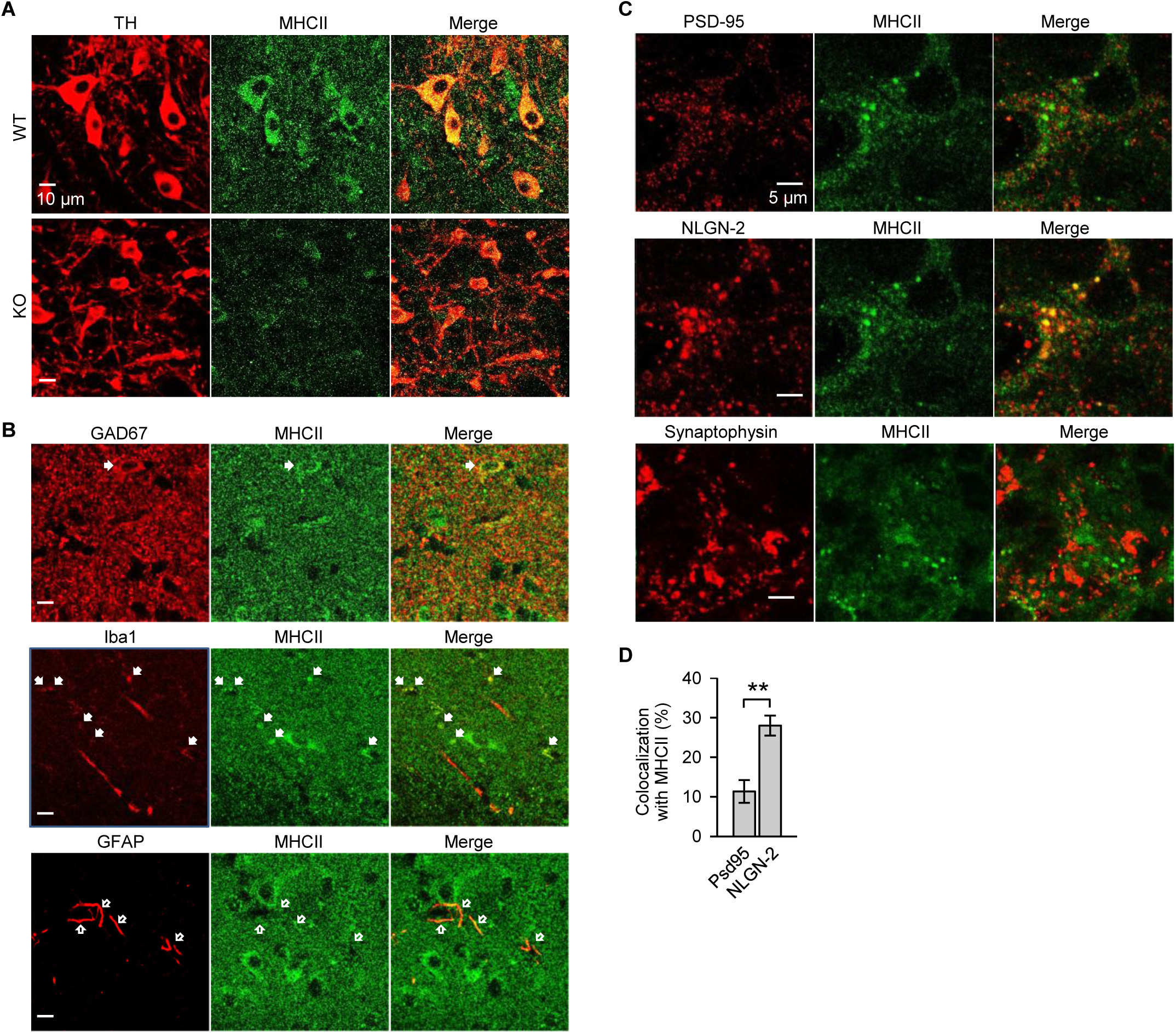
Immunohistochemical analysis of MHCII localization in the VTA. (A) Evaluation of antibody specificity against MHCII by double staining of MHCII with TH, a marker of dopaminergic neurons, in the VTA of WT and KO mice. (B) Identification of cells expressing MHCII by double staining of MHCII with GAD67, Iba1, and GFAP, a marker of GABAergic neurons, microglia, and glial cells in the VTA. (C) Identification of subcellular MHCII localization by double staining of MHCII expressed in the dopaminergic neurons with PSD-95, NLGN-2, and Synaptophysin, a marker of excitatory postsynapses, GABAergic postsynapses, and presynapses in the VTA. (D) Colocalization of MHCII with PSD-95 or NLGN-2 (*n* = 14) (Student’s t test, \*\**p* < 0.01). Data are represented as mean ± SEM.

Analysis of cell types expressing MHCII showed that soma-like staining of MHCII predominantly colocalized with tyrosine hydroxylase (TH), a marker of dopaminergic neurons. MHCII staining was also colocalized with glutamic acid decarboxylase (GAD65/67), and ionized calcium-binding adapter molecule 1 (Iba1), which are markers of GABAergic neurons and microglia, respectively (Figure 4B). In contrast, glial fibrillary acidic protein (GFAP), a marker of glial cells, did not overlap with MHCII. In the analysis of the subcellular localization, higher-magnification images revealed punctate MHCII immunostaining within and outside the soma of dopaminergic neurons. This staining overlapped extensively with the GABAergic postsynaptic marker neuroligin-2 (NLGN-2), slightly with the excitatory postsynaptic marker postsynaptic density protein-95 (PSD-95), and minimally with the presynaptic marker synaptophysin (Figures 4C and 4D). As dopaminergic neurons account for more than 70% of the total population of VTA neurons, these findings indicate a preferential MHCII localization at GABAergic postsynaptic sites in dopaminergic neurons.

### MIA treatment-induced reduction of GABAergic synapses on dopaminergic neurons

Given the role of MHCI in synaptic pruning, our results suggest the possibility that MHCII may prune GABAergic synapses. To directly confirm this hypothesis, we investigated GABAergic synaptic transmission to dopaminergic neurons by analyzing miniature inhibitory postsynaptic currents (mIPSCs) using ex vivo whole-cell patch-clamp recordings in dopaminergic neurons. Compared with saline-treated mice, MIA-treated mice exhibited lower mIPSC frequency but not a significant alteration in amplitude (Figures 5A-5C). This result suggests that MIA treatment reduced the number of GABAergic synapses. It is important to note that our immunohistochemical analysis revealed no difference in the number of GABAergic neurons between MIA- and saline-treated mice, indicating that the reduced number of GABAergic synapses in MIA-treated mice was not due to a decrease in GABAergic neurons (Figure 5D).

**Figure 5.**
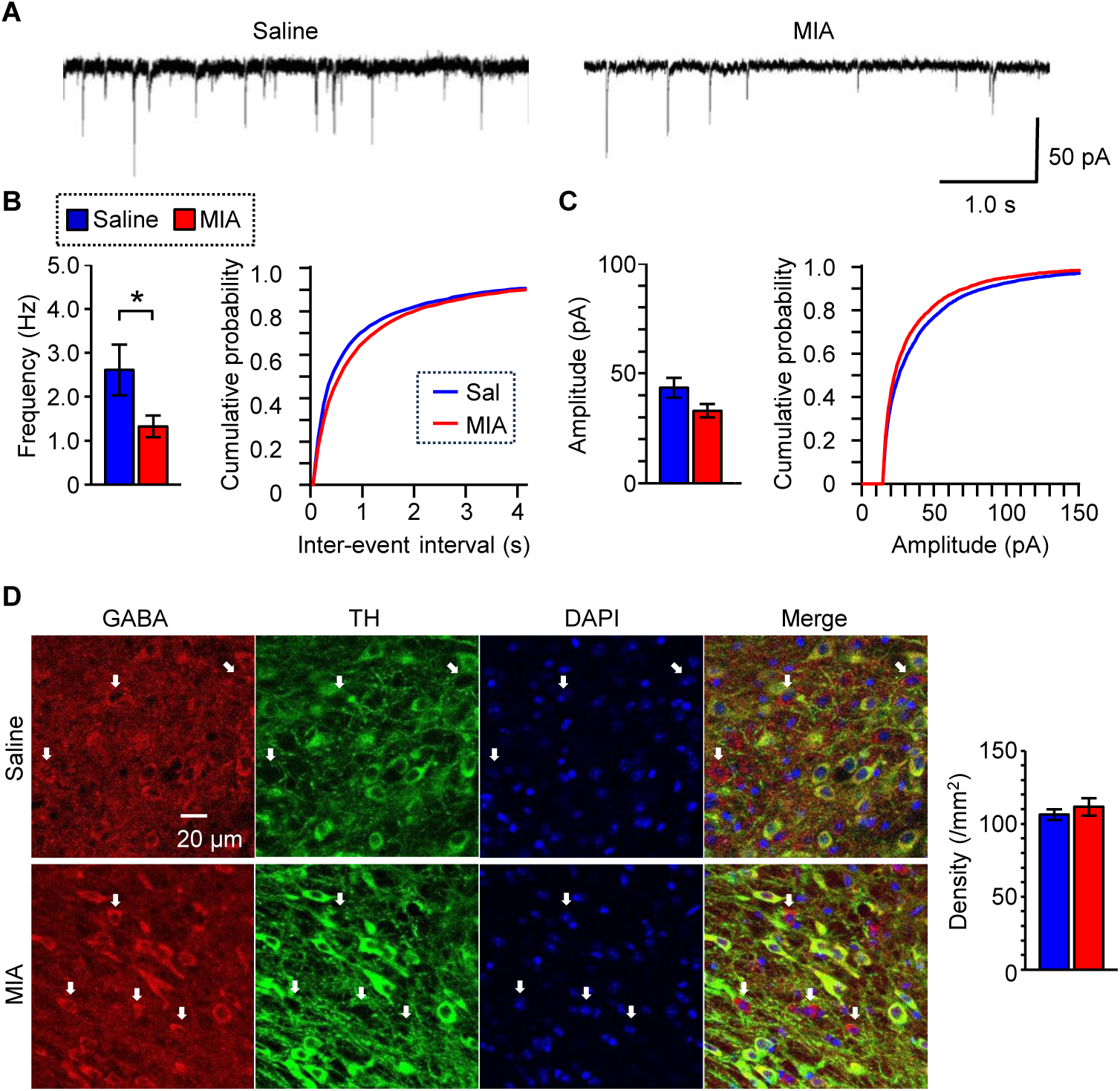
Ex vivo whole-cell patch-clamp recordings in VTA dopaminergic neurons. (A) Representative traces of mIPSCs in dopaminergic neurons of mice treated with saline or MIA. (B and C) Mean (left) and cumulative probability (right) of mIPSC frequency (B) and amplitude (C) in dopaminergic neurons of mice treated with saline (*n* = 17) or MIA (*n* = 19) (Student’s t-test, \**p* < 0.05). (D) Representative images of immune triple staining of GABA (red), TH (green), and DAPI (blue) for identification of GABAergic neurons (left), and density of GABAergic neurons (right) in the VTA of mice treated with saline (*n* = 6) or MIA (*n* = 6). Arrows indicate GABAergic neurons with staining of GABA and DAPI but not TH. Values are expressed as mean ± SEM and labeled as in (B).

### MHCII KO-induced enhancement of GABAergic synapses

Because our qPCR analysis showed negative correlations between MHCII and *Gad* expression, it is important to investigate the causal relationship between them. For this purpose, we examined the effect of MHCII KO on GABAergic synapses and behaviors using MHCII KO mice. qPCR analysis confirmed that deletion of classical MHCII expression induced an increase in Gad1/2 expression in the VTA of KO mice compared with WT mice (Figure 6A). This increase in *Gad* expression was also confirmed at the protein level in the VTA, as well as in the hippocampus, likely due to MHCII deletion throughout the brain in KO mice (Figures 6B and S3).

**Figure 6.**
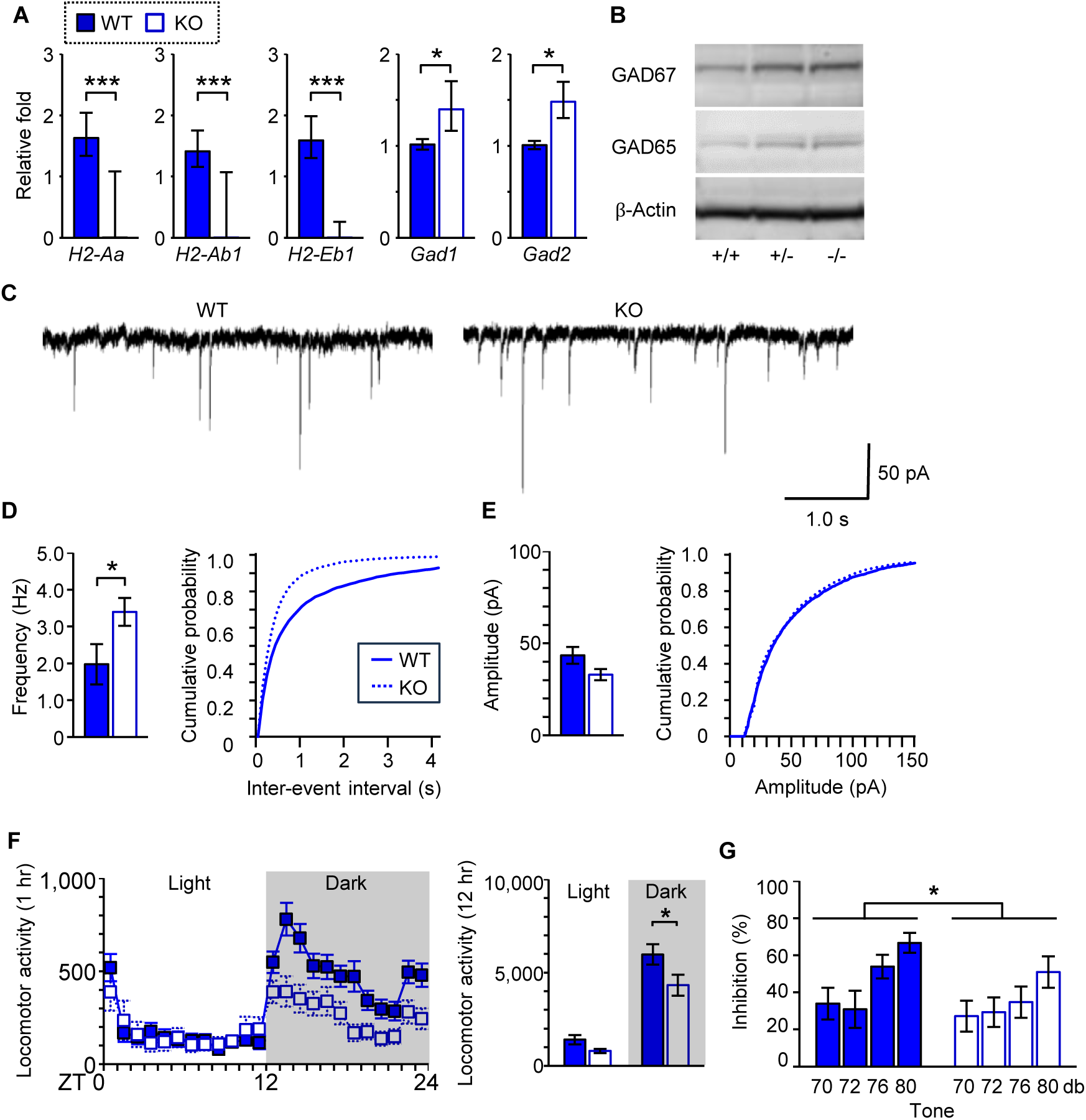
Contribution of MHCII KO on GABAergic synapses and behaviors. (A) MHCII and *Gad* mRNA expression in the VTA of WT (*n* = 11) and homo KO (*n* = 5) mice (Student’s t test, \**p* < 0.05, and \*\*\**p* < 0.001). (B) Expression of GAD protein in the VTA of WT (+/+), heterozygous KO (+/−), and homozygous KO (−/−) MHCII mice. (C) Representative traces of mIPSCs in dopaminergic neurons of WT and homo KO mice. (D and E) Mean (left) and cumulative probability (right) of mIPSC frequency (D) and amplitude (E) in dopaminergic neurons of WT (*n* = 13) and homo KO (*n* = 13) mice (Student’s t-test, \**p* < 0.05). (F) Locomotor activity of WT (*n* = 18) and homo KO (*n* = 13) mice (Student’s t-test, \**p* < 0.05). (G) PPI ratio calculated by the response to a startle stimulus (120 dB) with a weak pre-stimulus (70, 72, 76 and 80 dB) divided by the startle stimulus only in WT (*n* = 7) and homo KO (*n* = 10) mice (two-way ANOVA, stimulus: F_(3, 84)_ = 5.6, \*\**p* < 0.01; treatment: F_(1, 84)_ = 4.1, \**p* < 0.05; stimulus × treatment: F_(3, 84)_ = 0.69, p = 0.56).

Next, inhibitory synaptic transmission was analyzed using whole-cell patch-clamp recordings in dopaminergic neurons. MHCII KO mice exhibited increased mIPSC frequency with similar amplitude to WT mice (Figures 6C-6E). These mice also showed behavioral deficits, such as decreased locomotor activity and reduced PPI, similar to the behavioral disturbances seen in MIA-treated mice (Figures 6F and 6G). These results indicate that MHCII prunes GABAergic synapses and contributes to the observed behavioral deficits.

### Overexpression of MHCII in dopaminergic neurons

Because MHCII expression was deleted throughout the body in MHCII KO mice and our immunohistochemical analysis identified MHCII expression in dopaminergic neurons, as well as GABAergic neurons and microglia, it is important to specify which cell type’s expression of MHCII contributes to the pruning of GABAergic synapses in the VTA. For this purpose, we overexpressed *H2-Aa* specifically in dopaminergic neurons, dominant neurons in the VTA. We used rAAV-2 encoding *H2-Aa* conjugated with enhanced green fluorescent protein (*EGFP*), or *EGFP* only as a control, under the TH promoter (Figure 7A). Microinjection of rAAV-2 into the VTA increased *H2-Aa* expression specifically in dopaminergic neurons. This increase was accompanied by elevated expression of other classical MHCII including *H2-Ab1* and *H2-Eb*, which was consistent with the positive correlations among MHCII genes observed in our qPCR analysis. This overexpression of *H2-Aa* induced a decrease in Gad1/2 expression. Correlation analysis also revealed negative correlations between classical MHCII and *Gad1/2* expression (Figure 7D). These results indicate that MHCII expression in dopaminergic neurons is sufficient for pruning of GABAergic synapses.

**Figure 7.**
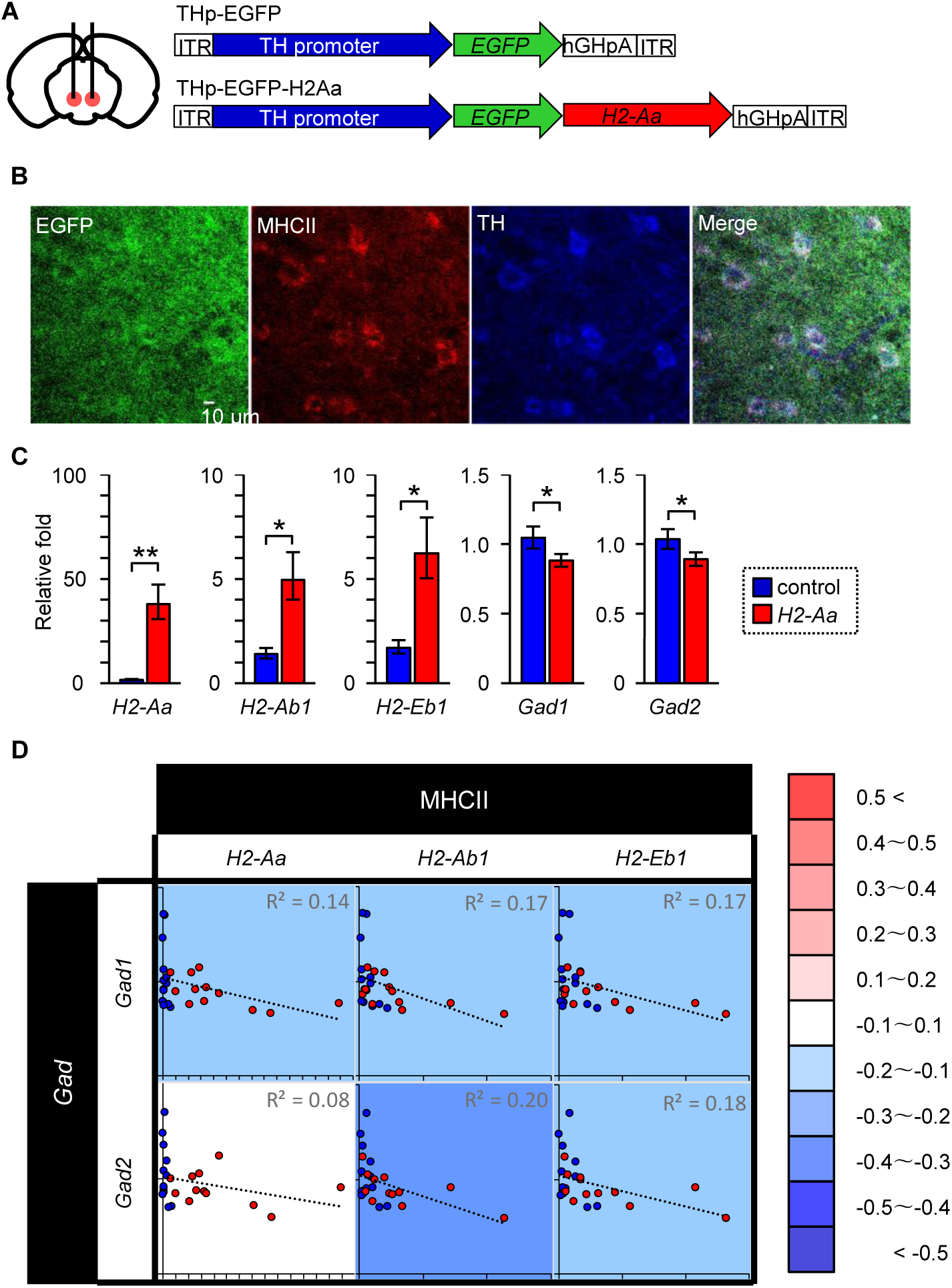
Contribution of *H2-Aa*/MHCII overexpression specifically in dopaminergic neurons on *Gad* expression. (A) Recombinant AAV-2 constructs encoding *H2-Aa* conjugated with *EGFP* or *EGFP* only under control of the TH promoter (THp). ITR, inverted terminal repeat; hGHpA, human growth hormone polyA sequence. (B) Representative images of triple immunostaining for EGFP (green), MHCII (red), and TH (blue), demonstrating MHCII overexpression specifically in dopaminergic neurons. (C) MHCII and *Gad* mRNA expression in the VTA of control (*n* = 14) and *H2-Aa* overexpressing (*n* = 12) mice (Student’s t test, \**p* < 0.05, and \*\**p* < 0.01). (D) Correlation analysis of MHC and *Gad* mRNA expression in the VTA of control (*n* = 12) and *H2-Aa* overexpressing (*n* = 9) mice. *p* and *r*^2^values are shown in the upper left and upper right corners respectively. All data are represented as mean ± SEM and labeled as in (C).

## Discussion

The results of the present study suggest that MIA treatment increases classical MHCII expression at postsynaptic sites of GABAergic synapses in dopaminergic neurons and enhances the pruning of GABAergic synapses on these neurons. These alterations persist into adulthood, contributing to behavioral deficits characteristic of neurodevelopmental disorders. Our behavioral analyses demonstrated that MIA treatment induced behavioral deficits in locomotor activities and PPI, which are characteristically impaired in ADHD and schizophrenia, respectively. Although autism- and cocaine addiction-associated behavioral abnormalities were not observed, these analyses focused on specific aspects of behavioral deficits; broader investigations are necessary for a comprehensive evaluation.

### Alterations in gene expression by MIA treatment

Neurodevelopmental disorders are polygenic, and our RNA-seq analysis also identified genes associated with these disorders other than MHCII and GABAergic synapse-related genes such as *Drd5* and *Syn1* particularly in the VTA. *Drd5* belongs to the D1-like receptor family, and a significant association between ADHD and a microsatellite marker located near the gene has been revealed by a joint analysis of 14 independent samples ^22^. *Syn1* plays a crucial role in neurotransmitter release and neuronal plasticity, and variants in the *gene* have been associated with X-linked neurodevelopmental disorders ^23^. Because *Drd5* and *Syn1* are associated with GABAergic synapses, alterations in the expression of these genes are consistent with our findings showing MHCII-mediated pruning of GABAergic synapses. Although the mechanism of VTA-specific gene alterations in MIA-treated mice was not addressed in this study, dopaminergic neurons have been reported to be more sensitive to cytokines ^16^. In our study, MIA treatment induced proinflammatory cytokine expression in the fetal whole-brain within 3 hours. Among these cytokines, *Ifng* is recognized as the most prominent regulator of MHCII expression; thus, it may contribute to the persistent upregulation of MHCII expression in the VTA by MIA treatment.

### MHCII expression in other cells than dopaminergic neurons

Regarding MHCII expression in the brain, many studies have focused on MHCII expressed in microglia; however, other studies have demonstrated MHCII expression in neurons, including dopaminergic neurons ^24–26^. Furthermore, single-cell RNA-seq studies have revealed MHCII expression in both dopaminergic neurons and microglia ^27,28^. Our study also showed MHCII expression in dopaminergic neurons, GABAergic neurons, and microglia, and MHCII expression in dopaminergic neurons was sufficient for pruning of GABAergic synapses. Although its expression level is considered low, our previous studies have demonstrated that genes with low expression can play important roles in synaptic microstructure ^21^. As our study did not address the contributions of MHCII expressed in GABAergic neurons or microglia to pruning of GABAergic synapses, it is important to investigate these contributions in future study.

### GABAergic synapse in neurodevelopmental disorders

Disruption of GABAergic synapses is a primary factor contributing to neurodevelopmental disorders, including autism spectrum disorder and schizophrenia. ^29^. MIA treatment has also been shown to affect multiple steps of cortical GABAergic interneuron development, such as proliferation of precursor cells, migration, and positioning of neuroblasts, and neuronal maturation ^30^. Furthermore, increased cytokine levels in the fetal brain have been reported to compromise the migration of cortical GABAergic neurons ^31,32^. In contrast to these studies, our study demonstrated that MIA treatment disrupted GABAergic synapses through the regulation of MHCII in dopaminergic neurons, rather than GABAergic neurons themselves. Therefore, our findings reveal a distinct mechanism compared to the disruption of GABAergic neurons. Notably, GABAergic synapses are the primary inhibitory synapses in the central nervous system, while glycinergic synapses, another type of inhibitory synapse, are not significantly present or functional in the VTA ^33^.

### Type 1 diabetes like association between MHCII and GAD

Although our study suggested a novel mechanism of neurodevelopmental disorders whereby MHCII prunes GABAergic synapses, a similar mechanism has been observed in type 1 diabetes. In this disease, GAD peptides originating from pancreatic β-cells are recognized by MHCII in antigen-presenting cells, and presentation of these peptides to T cells triggers activation of an autoimmune reaction against β-cells, resulting in β-cell death . Applying this mechanism to our results, it is possible that GAD peptides exposed from presynaptic sites of GABAergic synapses are recognized by MHCII at the postsynaptic sites, and presentation of these peptides triggers pruning of GABAergic synapses (Figure S4). Because T cells are absent or scarce in the healthy brain parenchyma, microglia might assume T cell-like roles in this context. Although this possibility should be addressed in future study, meta-analysis has shown a higher prevalence of autism spectrum disorders and ADHD in the type 1 diabetes population, suggesting a common mechanism underlying these conditions ^34^.

Given that dopaminergic neurons are implicated in various psychiatric and neurological disorders, it is possible that disturbances in MHCII expression may also underlie these conditions. Indeed, progressive loss of dopaminergic neurons is a hallmark of Parkinson’s disease, and growing evidence suggests that MHCII plays an important role in its pathogenesis ^25^. Furthermore, MHCII is expressed in various types of neurons and implicated in some neurological diseases such as Multiple Sclerosis or Narcolepsy ^35,36^. Therefore, our finding that MHCII mediates pruning of GABAergic synapses might contribute to elucidating the pathogenesis of various psychiatric and neurological disorders.

## Acknowledgments

We are grateful to members of the Department of Biochemistry and members of the Biomedical Research Center, Saitama Medical University, for their help with this work. The animals used in this study were maintained with the help of the staff of the Division of Experimental Animals, Saitama Medical University. Funding: This study was supported by the Grants-in-Aid for Scientific Research (grant nos. JP 25K13174, JP24K10686, JP20K07089, and JP17K08960) from the Japan Society for the Promotion of Science (to G.M.); Author Contributions: G.M. designed and performed all experiments, analyzed data, and wrote the manuscript. M.Hi. performed RNA-seq experiments. M.Ha. assisted in whole-cell patch-clamp recordings. T.M. reviewed results for experiments. Data and materials availability: All data needed to evaluate the conclusions are presented in the paper and/or Supplementary Materials. Additional data related to this paper may be requested from the authors.

## Material and Methods

### Animals and MIA treatment

For MIA treatment and overexpression of *H2-Aa* in dopaminergic neurons, C57BL/6J mice were purchased from Japan SLC (Hamamatsu, Japan) and Jackson Laboratory Japan (Yokohama, Japan). In MIA treatment, Poly(I:C) (MilliporeSigma, Burlington, MA, USA) was intraperitoneally injected at a dose of 5.0 mg/kg body weight into pregnant mice on gestation day 9. Knockout (KO) mice with a constitutive homozygous deletion of all classical MHCII (*H2^dlAb^*^1^*^-Ea^*) were purchased from the Jackson Laboratory (Bar Harbor, ME, USA), and maintained as heterozygous mutants on a C57BL6/J background. For comparative analysis of KO and wild-type (WT) mice, both genotypes were generated by breeding heterozygous mutants. All mice were individually housed after weaning at 4 weeks of age under a standard 12-hour light/12-hour dark cycle (lights on at 07:00), at a controlled temperature of 25°C and a relative humidity of 60%, with *ad libitum* access to food and water. All procedures were approved by the Animal Care and Use Committee of Saitama Medical University and were conducted in accordance with institutional guidelines. All efforts were made to minimize both the suffering and the number of animals used.

### Locomotor activity

After acclimation to an open-field test box (30 × 30 × 35 cm) for at least 3 hours, locomotor activity was measured over an average period of more than three consecutive days under a regular 12-hour light/12-hour dark cycle, with *ad libitum* access to food and water. The apparatus was equipped with infrared sensors attached to the lid (Biotec Co., Ltd., Osaka, Japan) and clean paper bedding on the floor.

### Prepulse inhibition (PPI)

Mice were placed in a startle chamber (Startle Reflex. Test Unit; O’Hara & Co., Ltd, Tokyo, Japan) for 10 min with a baseline background noise of 70 dB for habituation. The background noise continued throughout the experiment. Subsequently, the mice were subjected to ten trials of a 40-ms, 120-dB startle stimulus. This was followed by twenty trials presented in pseudo-random order, consisting of either no stimulus or a 20-ms prepulse stimulus of 70, 72, 76, or 80 dB, each followed by a 40-ms, 120-dB startle stimulus 100 ms later. Following this, ten trials of 40 ms, 120-dB startle stimulus were presented again. The interval between trials ranged from 20 to 30 seconds. Startle responses were recorded after the startle stimulus and the peak values were analyzed. Percent PPI was calculated as: 100 − {[(startle response for startle stimulus with prepulse stimulus) / (startle response for startle stimulus alone)] × 100}.

### Social behaviors

Social interaction and social recognition were assessed using the three-chamber test. The apparatus consisted of a rectangular box (42 × 60 × 22 cm) divided equally into three chambers by Plexiglas partitions with small rectangular openings. An empty wire cylinder was placed in each of the side chambers. For habituation, the test mouse was introduced into the center chamber and allowed to explore all three chambers, including the empty cylinders, for 10 min. In the subsequent 10-min session, an unfamiliar conspecific male mouse (Stranger 1) of similar weight was placed in one of the empty cylinders. Social interaction was quantified as the difference between time spent near the cylinder containing Stranger 1 and time spent near the empty cylinder. In the final 10-min session, a novel conspecific male mouse (Stranger 2) of similar weight was placed in the previously empty cylinder. Social recognition was assessed by calculating the difference between time spent near Stranger 2 and time spent near Stranger 1.

### Cocaine-induced behavioral sensitization

After acclimation to an open-field test box and intraperitoneal saline injections over 4 days, mice were subjected to seven daily injections of cocaine at doses of 1.0 or 2.0 mg/kg body weight. Following a 14-day withdrawal period, all groups received a challenge injection of cocaine at the same dose. Locomotor activity was recorded throughout the experiment.

### RNA-seq

Brain regions were dissected from three saline-treated and three MIA-treated mice. The regions of interest included the ventral tegmental area (VTA), nucleus accumbens (NAc), medial prefrontal cortex (mPFC), and hippocampus (Hip). These brain regions were excised, homogenized, and subjected to RNA extraction as described in the qPCR analysis section. The RNA library was prepared from high-quality RNA depleted of rRNA. After adaptor ligation, the resulting DNA was amplified by PCR for 12 cycles and purified. Resulting libraries were quantified and then used for cluster generation and sequencing. For analysis of RNA-sequence data, low-quality bases were removed from the reads and the resulting trimmed sequencing reads were aggregated into a rRNA reference to remove rRNA reads. Then, the clean reads were mapped to the grcm38_snp_tran reference genome and sorted. Gene expression levels were measured using TPM (transcripts per million) and calculated using R package Ballgown. The readcounts were calculated using HTseq. *p* values for the difference between groups were obtained using the edgeR package (https://bioconductor.org/packages/release/bioc/html/edgeR.html).

### qPCR

Mice were deeply anesthetized and decapitated. Brains were rapidly removed and frozen on dry ice and stored at −80°C until slicing. Coronal brain slices (1-mm thickness) were prepared using a mouse brain matrix. Then, brain regions including the VTA, NAc, mPFC, and Hip were dissected using a razor blade. Total RNA was isolated from each region using TRI Reagent (Funakoshi, Tokyo, Japan). cDNA was synthesized from total RNA using a High-Capacity cDNA Archive Kit (Thermo Fisher Scientific, Waltham, MA, USA). Quantitative real-time PCR was performed using the QuantStudio 12K Flex (Thermo Fisher Scientific). All primers were from TaqMan Gene Expression Assays (Thermo Fisher Scientific) and are listed in Table 1.

### Western blot

Brain regions were prepared as described in the qPCR section. The brain regions were homogenized in lysis buffer containing 1% SDS, 1% TritonX-100, and protease inhibitor cocktail (EZBlock; Biovision Inc., Mountain View, CA). Then, the protein concentration was quantified using a BCA kit (Thermo Fisher Scientific), and samples were prepared at a concentration of 1.0 mg/mL using sample buffer (AE-1430CP EzApply; ATTO, Tokyo, Japan). Twenty micrograms of each sample were subjected to electrophoresis on a 15% precast gel (E-R15L; ATTO) and transferred to polyvinylidene fluoride membranes (Clear Blot Membrane-p AE-6665; ATTO) using blotting buffer (AE-1465 EzFastBlot; ATTO). The membranes were incubated with blocking buffer containing phosphate-buffered saline (PBS), 3% skim milk (Wako, Tokyo, Japan), and 0.1% Tween-20 (Wako) or with Pierce Protein-Free Blocking Buffer (Thermo Fisher Scientific) for 15 minutes at room temperature. Subsequently, the membranes were incubated with primary antibodies overnight at 4°C, followed by incubation with secondary antibodies in Pierce Protein-Free Blocking Buffer for 1 hour at room temperature. Protein bands were detected using enhanced chemiluminescence (ECL) plus western blotting detection reagents (Cytiva, Marlborough, MA, USA). Images of chemiluminescence from protein bands were obtained using a ChemiDoc Touch MP imaging system (Bio-Rad, Hercules, CA, USA). The primary antibodies and their dilutions are listed in Table 2.

### Immunohistochemistry

Mice were deeply anesthetized and transcardially perfused with PBS for 5 min, followed by PBS containing 4% paraformaldehyde for 5 min. Whole brains were dissected, and 40-μm-thick slices including the targeted brain regions were prepared on a cryostat (M1950; Leica Biosystems, Wetzlar, Germany). Brain slices were incubated for 30 minutes at room temperature in blocking buffer containing PBS, 5% normal goat serum (Wako), and 10% bovine serum albumin (MilliporeSigma), and 0.5% Triton X-100. Slices were then incubated overnight at 4°C with primary antibodies, followed by incubation with secondary antibodies for 30 minutes at room temperature in a staining buffer prepared by diluting the blocking buffer to one-third of its original concentration. Images were captured using confocal microscopy (Axio Imager Z2, LSM710 CLSM; Zeiss, Oberkochen, Germany). The primary antibodies and their dilutions are listed in Table 2. The percent colocalization with MHCII was calculated as: [(colocalized MHCII intensity) / (total MHCII intensity)] × 100.

### Whole-cell patch-clamp recordings

Mice were deeply anesthetized and transcardially perfused with 25 mL ice-cold sucrose solution containing (in mM): 220 sucrose, 2.5 KCl, 1.25 NaH_2_PO_4_, 10.0 MgSO_4_, 0.5 CaCl_2_, 26.0 NaHCO_3_, 30.0 glucose, 3.0 sodium pyruvate, and 1.0 sodium ascorbate. Brains were rapidly removed and submerged in cold oxygenated sucrose solution for 1 min. After removing the forebrain, cerebellum, and dorsal portion of the brain with razor blades, horizontal brain slices (300 µm thick) containing the VTA were cut in sucrose solution using a microslicer (VT1000S; Leica Microsystems AG) with a ceramic blade (7550/1/C; Campden Instruments Limited, Loughborough, UK). Slices were placed in a custom-made interface storage chamber containing artificial cerebrospinal fluid (ACSF) consisting of (in mM): 125 NaCl, 2.5 KCl, 1.25 NaH_2_PO_4_, 1.0 MgSO_4,_ 2.0 CaCl_2_, 26.0 NaHCO_3_, 1.0 ascorbic acid-Na, and 20.0 glucose, aerated with 95% O_2_ and 5% CO_2_ at 33°C for 2 hours. Slices were maintained at room temperature until recordings.

Patch electrodes were fabricated from standard glass capillaries (1B150F-4; World Precision Instruments, Sarasota, FL, USA) using a P-97 horizontal puller (Sutter Instruments, Novato, CA, USA). Electrode resistance ranged from 3–6 MΩ when filled with an internal solution containing (in mM): 150 CsCl, 5 KCl, 0.1 CsEGTA, 5 Cs-HEPES, 3 Mg-ATP, and 0.4 Na_2_GTP. For cell type identification, the pipette solution also included 0.2% biocytin. Individual slices were transferred to a recording chamber, perfused at 2 mL/min with aerated ACSF, and maintained at 32°C. Dopaminergic neurons in the VTA were identified by the presence of large hyperpolarization-activated currents during hyperpolarizing pulses from −70 mV. For recordings of miniature inhibitory postsynaptic currents (mIPSCs), neurons were voltage-clamped at −65 mV in the presence of 10 μM 6-Cyano-7-nitroquinoxaline-2,3-dione (CNQX), 50 μM D-AP5, and 0.5 μM tetrodotoxin (TTX), and spontaneous activity was recorded for an average of 10 min. Recorded currents were acquired at 5 kHz and filtered at 1 kHz using a DigiData1322A and pClamp 9 software (Molecular Devices, Sunnyvale, CA, USA). Data were analyzed using Clampfit 9 (Molecular Devices).

### Microinjection of rAAV-2 in VTA

Mice were deeply anesthetized and placed in a stereotaxic instrument (Narishige). A hole was made on each side of the skull with a dental drill, and a Hamilton microsyringe with 32-gauge needle (Hamilton, Reno, NV, USA) vertically inserted into the VTA (2.7 mm posterior, 0.5 mm lateral, and 4.0 mm ventral to bregma). A 0.4 μL volume of recombinant adeno-associated virus (rAAV) (titer 6.0 × 10^11) encoding *H2-Aa* conjugated with *EGFP* or *EGFP* only under control of the TH promoter was microinjected into each side for 5 min. The needle was kept in place for an additional 5 min before it was slowly withdrawn. Thirty days after the surgery, qPCR analysis was conducted. The rAAV vector was purchased from Vector Builder (IL, USA).

### Statistical analyses

All data are expressed as mean ± SEM. All statistical analyses were performed with IBM SPSS software (IBM Corp., Armonk, NY, USA). Unpaired two-tailed Student’s t-tests were applied to identify significant differences between two groups after verifying the normality assumptions. Statistical significance for multiple group comparisons was primarily assessed using analysis of variance (ANOVA), followed by post hoc tests when applicable. For correlation analysis, the following equation: *t_0_* = *r* / sqrt[(1 − *r*^2^) / (*n* − 2)] was used. In this equation, *r* represents the correlation coefficient, *n* the number of samples, and then *t_0_* distributes as *t* with *d.f*. = *n* − 2.

---

**Figure S1.**
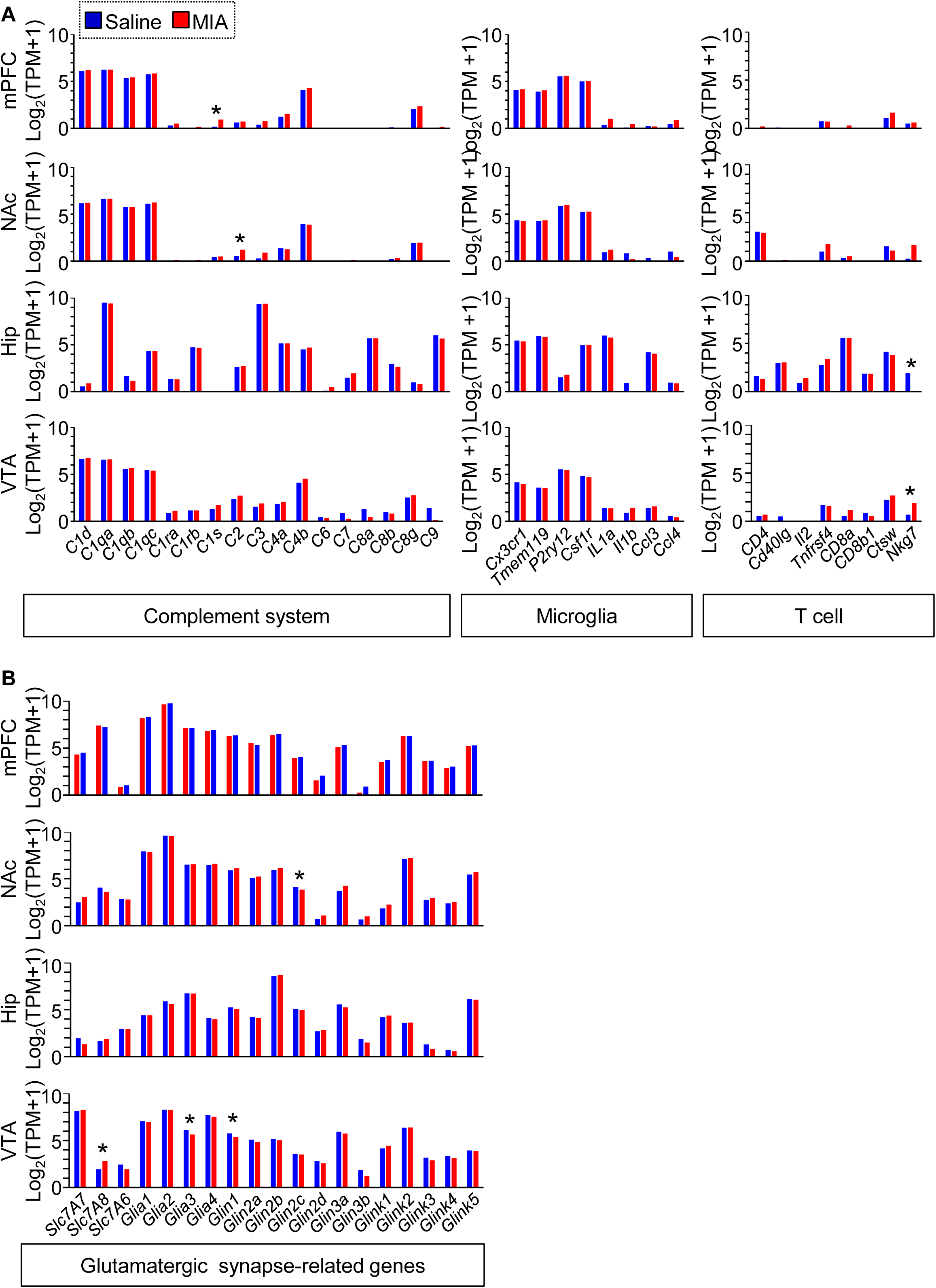
MIA treatment-induced alterations in immune-related and glutamatergic synapse-related gene expression. RNA-seq analysis of genes associated with the complement system, microglia, and T cells (A) and glutamatergic synapses (B) in the VTA, mPFC, NAc, and Hip of mice treated with saline (*n* = 3) or MIA (*n* = 3). All data are represented as mean ± SEM. \**p* < 0.05

**Figure S2.**
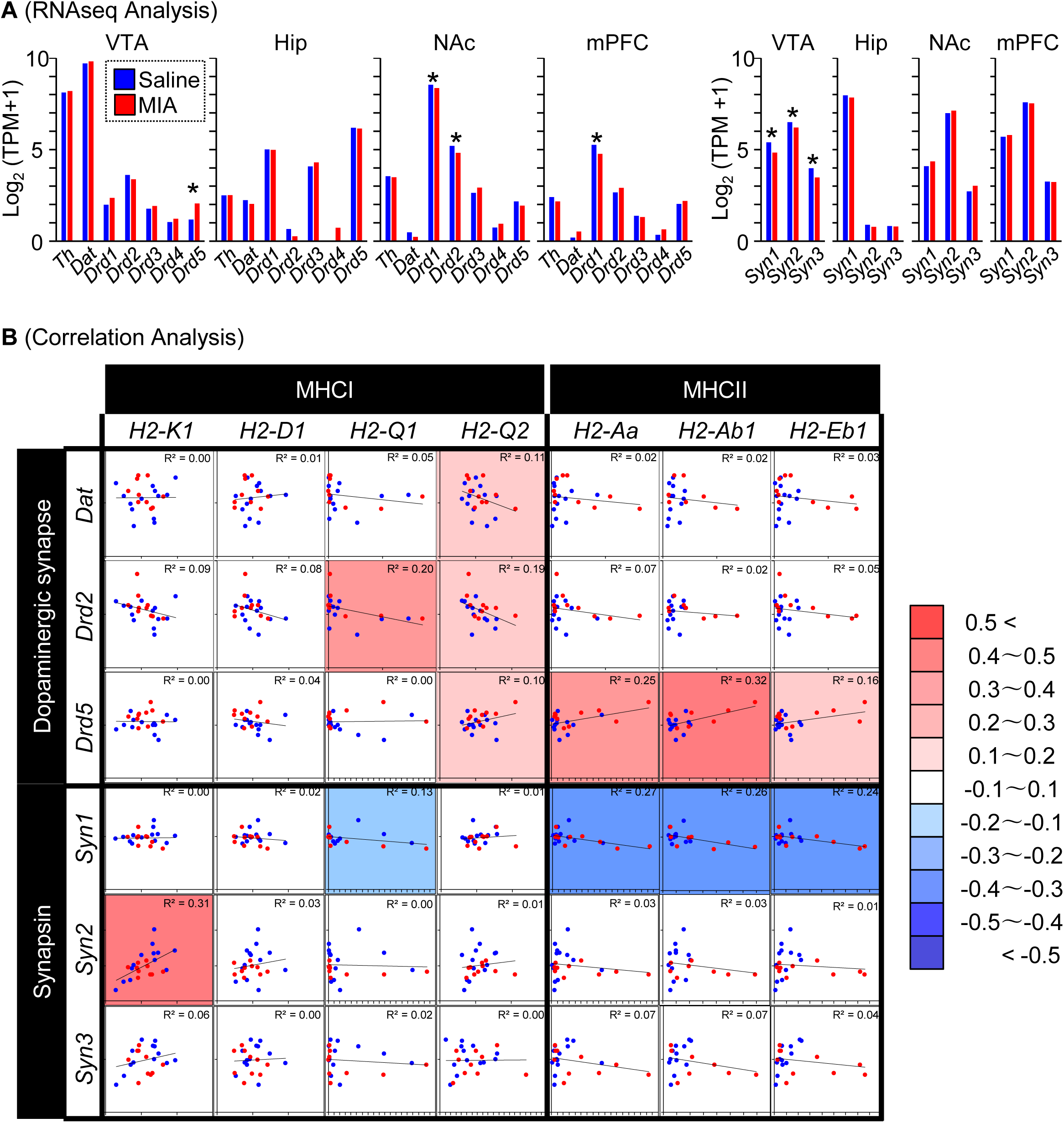
MIA treatment-induced alterations in dopaminergic neuron-related gene and synapsin expression. (A) RNA-seq analysis of dopaminergic neuron-related gene and synapsin expression in the VTA, mPFC, NAc, and Hip of mice treated with saline (*n* = 3) or MIA (*n* = 3) (\**p* < 0.05, \*\*\**p* < 0.001). (B) Correlation analysis of MHC with dopaminergic neuron-related gene and synapsin expression in the VTA of mice treated with saline (*n* = 13) or MIA (*n* = 10). *r*^2^values are shown in the upper right corners. All data are represented as mean ± SEM and labeled as in (A).

**Figure S3.**
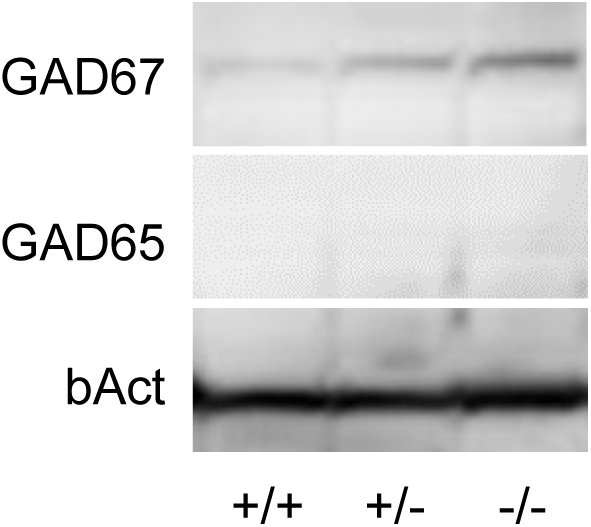
Expression of GAD protein in the Hip of WT and KO mice. Expression of GAD protein in the Hip of WT (+/+), heterozygous KO (+/−), and homozygous KO (−/−) MHCII mice.

**Figure S4.**
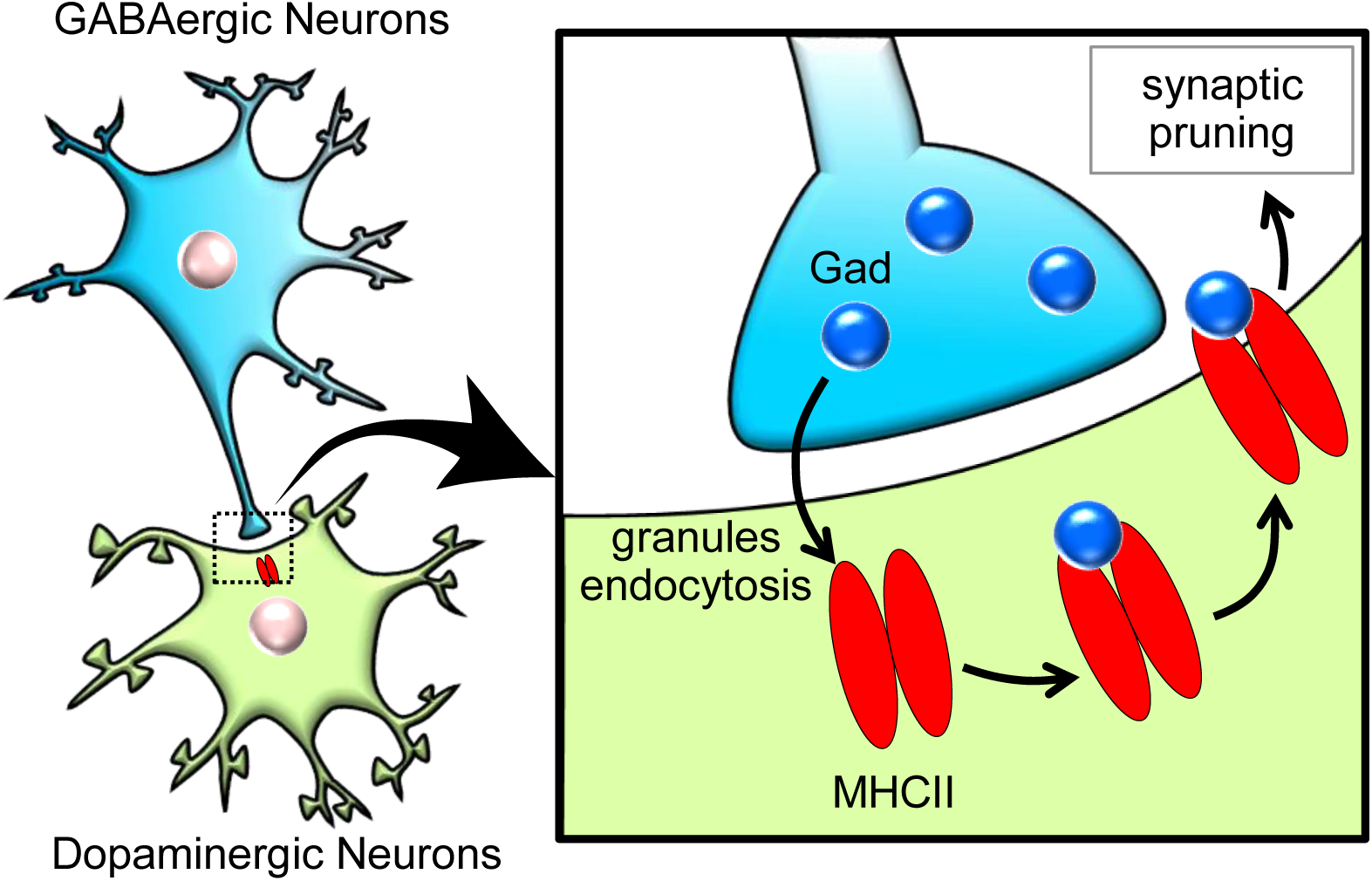
Schematic diagram of MHCII-mediated pruning of GABAergic synapses on dopaminergic neurons. GAD peptides exposed from presynaptic sites of GABAergic synapses are recognized by MHCII at the postsynaptic sites, and presentation of these peptides triggers pruning of GABAergic synapses.

